# Edaravone-Loaded Mesoscale Nanoparticles Treat Cisplatin-Induced Acute Kidney Injury

**DOI:** 10.1101/2020.01.24.919134

**Authors:** Ryan M. Williams, Janki Shah, Elizabeth Mercer, Helen S. Tian, Justin M. Cheung, Madeline Dorso, Edgar A. Jaimes, Daniel A. Heller

**Affiliations:** Memorial Sloan Kettering Cancer Center, New York, NY 10065; City College of New York Department of Biomedical Engineering, New York, NY 10031; Weill Cornell Medical College, New York, NY 10065

## Abstract

Cisplatin-induced acute kidney injury (CI-AKI) is a significant co-morbidity of chemotherapeutic regimens. While this condition is associated with substantially lower survival and increased economic burden, there is no pharmacological agent to effectively treat CI-AKI. The disease is hallmarked by acute tubular necrosis of the proximal tubular epithelial cells primarily due to increased oxidative stress. In our prior work, we developed a highly-selective kidney-targeted mesoscale nanoparticle (MNP) that accumulates primarily in the renal proximal tubular epithelial cells while exhibiting no toxicity. Here, we found that MNPs exhibit renal-selective targeting in multiple mouse models of tumor growth with virtually no tumor accumulation. We then evaluated the therapeutic efficacy of MNPs loaded with the reactive oxygen species scavenger edaravone in a mouse model of CI-AKI. We found a marked and significant therapeutic effect with this approach as compared to free drug or empty control MNPs, including improved renal function, histology, and diminution of oxidative stress. These results indicated that renal-selective MNP edaravone delivery holds substantial potential in the treatment of acute kidney injury among patients undergoing cisplatin-based chemotherapy.

## Introduction

Acute kidney injury (AKI) is a common clinical condition associated with significant morbidity and mortality regardless of etiology or setting. AKI affects millions of individual patients and has a large socioeconomic impact, including longer hospital stay and higher costs. In the US alone, it is estimated that the annual costs related to AKI are up to $24 billion^1^. The incidence of AKI is increasing at a rapid pace^2,3^, which is attributable to several factors including shifts in demographics, severity of underlying diseases, and expansion of invasive and complex medical procedures^4,5^. AKI can result from a variety of insults including volume depletion, septicemia, hypotension, and commonly used drugs including antibiotics and chemotherapeutic agents.

Cisplatin is a widely used chemotherapy in the treatment of a variety of cancers including ovarian, head and neck, bladder, testicular, and lung, among others^6,7^. A significant side effect of cisplatin therapy is the occurrence of AKI in approximately 33% of patients^8,9^. Indeed, it has been reported that about 20% of all AKI incidences are caused by cisplatin^7,10^. The development of AKI in these patients can result in the interruption of chemotherapy or a change to less effective chemotherapies^8,9^. There is therefore a critical need to develop novel strategies to prevent or treat AKI induced by cisplatin. This would also have a direct impact on the oncologic outcomes of these patients whose treatment otherwise cannot be completed, or other less-effective chemotherapeutic agents must be used^11^.

There has been significant overall progress in our understanding of the epidemiology, pathophysiology and outcomes of AKI. Despite such substantial advances, almost no meaningful progress has been made in the treatment of AKI and there are no effective drugs currently approved for the treatment of AKI^12^. Hence, clinicians rely on conservative measures and renal replacement therapy if indicated for the management of AKI^13^. Numerous strategies have been proposed to prevent, ameliorate, or treat AKI. These include inhibition of inflammatory mediators, enhancement of renal perfusion by blocking vasoconstrictor mechanisms and/or enhancing vasodilation, attenuation of leukocyte infiltration, inhibition of the coagulation cascade, inhibition of reactive oxygen species (ROS), and administration of growth factors to accelerate renal recovery^14,15^. Most of these attempts have shown moderate success in attenuating AKI in animal models but have subsequently failed in clinical trials. The reasons for these failures are multifactorial but often include poor delivery of therapeutic agents to the proximal tubules, systemic toxicity at doses required to have a therapeutic effect, concomitant co-morbid conditions, suboptimal clinical trial design, and heterogeneity in cause and timing of AKI, among others^16^.

In prior work, we developed a mesoscale nanoparticle (MNP)^17,18^ biodegradable drug carrier that localizes with up to 26-fold greater efficiency to the kidneys than any other organ. The nanoparticles specifically target the renal cortex tubular epithelium, with about 2/3 being proximal tubular epithelial cell localization and about 1/3 being distal tubular epithelial cell localization. These particles are biodegradable and safe, degrading in the tubular epithelial cells and releasing their cargo over days to weeks.

No clinical methods currently exist to target the majority of a therapeutic specifically to the site of CI-AKI^19,20^. Most attempted therapeutic strategies for CI-AKI have been hindered by side effects and/or poor drug accumulation at the site of injury, in addition to poor timing of AKI diagnosis and therapy initiation^21-23^. If strategies existed to change drug pharmacokinetics to localize therapies to the proximal renal tubules, therapeutic efficacy would likely be significantly improved^24^.

Here, we investigated the potential clinical utility of these particles by first demonstrating that they localize specifically to the kidneys in mouse tumor models, mitigating potential off-target effects of targeted therapy. We next investigated the therapeutic efficacy of drug-loaded MNPs in a mouse model of CI-AKI when loaded with a reactive oxygen species scavenger. Indeed, oxidative stress is a predominant mechanism of injury in cisplatin-induced AKI^8,25^. We used the slightly hydrophobic scavenger edaravone, which is approved in the U.S. for ALS and in Japan for ischemic stroke and ALS^26,27^. We found that edaravone-loaded MNPs imparted substantial protection against CI-AKI as measured by renal function, histology, and oxidative stress levels. We expect that the successful translation of such a technology may have a major impact on the morbidity and mortality associated with CI-AKI and become a new paradigm in the prevention and/or treatment of renal diseases associated with chemotherapeutic regimens.

## Results and Discussion

### MNP Formulation and Characterization

To specifically deliver the free radical scavenger edaravone to the site of CI-AKI, we formulated kidney-targeted mesoscale nanoparticles to encapsulate the drug. Via nanoprecipitation as previously described^17,18^ of edaravone with the block-copolymer poly(lactic-co-glycolic) acid (PLGA) conjugated to acid-terminated polyethylene glycol (PEG), or PLGA-PEG, we obtained therapeutic MNPs (Eda-MNPs). We also obtained fluorescent MNPs for imaging via nanoprecipitation with the Cy-5 mimic fluorescent dye 3,3’-diethylthiadicarbocyanine iodide (DEDC-MNPs). Finally, we produced control MNPs with no encapsulated molecules via essentially the same procedure, apart from a lack of co-precipitate small molecule (Ctrl-MNPs).

To characterize MNPs, we first performed hydrodynamic diameter and ζ-potential assays (Table 1). Eda-MNPs exhibited an average diameter (z average) of 374.0 ± 12.2 nm with a polydispersity index (PDI) of 0.337. 85% of these particles were within the 164 – 531 nm size range (Table 1; Figure 1a). Eda-MNPs exhibited a ζ-potential of −22.8 ± 0.55 mV in distilled water and −18.2 ± 0.51 mV after incubation with 100% serum (Figure 1b). DEDC-MNPs exhibited an average diameter of 366.4 ± 1.85 nm with a PDI of 0.308. 94% of these particles fell within the range of 164 – 531 nm (Table 1; Figure 1a). DEDC-MNPs exhibited a ζ-potential of −25.6 ± 1.13 mV in distilled water and −15.1 ± 0.44 mV in 100% serum (Figure 1b). Finally, Ctrl-MNPs exhibited an average size of 332.0 ± 7.56 nm with a PDI of 0.302. 74% of these particles fell within the 164 – 531 nm size range (Table 1; Figure 1a). Ctrl-MNPs exhibited a ζ-potential in distilled water of −24.7 ± 0.61 mV and −13.2 ± 0.7 mV in 100% serum (Figure 1b). In two prior studies with MNPs, we found that particles of this composition with hydrodynamic diameters of 347.6 nm and 386.7 nm, PDIs of approximately 0.3, and surface charges in water of approximately −20 mV each had significant renal accumulation^17,18^. We also found that the negative surface potential was partially abrogated after serum incubation^17,18^. Therefore, we would expect that given the similar size and charge characteristics, as well as their interaction with serum, each of the formulations would exhibit similar renal targeting behavior.

**Table 1.** Physical characteristics of MNPs used in all studies.

| Formulation | Hydrodynamic Diameter (nm) | PDI | % within 164 – 531 nm | ζ-potential in distilled water (mV) | ζ-potential in 100% serum (mV) |
| --- | --- | --- | --- | --- | --- |
| Eda-MNPs | 374.0 (12.2) | 0.337 (0.032) | 85% | -22.8 (0.55) | -18.2 (0.51) |
| DEDC-MNPs | 366.4 (1.85) | 0.308 (0.041) | 94% | -25.6 (1.13) | -15.1 (0.44) |
| Ctrl-MNPs | 332.0 (7.56) | 0.302 (0.040) | 74% | -24.7 (0.61) | -13.2 (0.70) |

**Figure 1.**
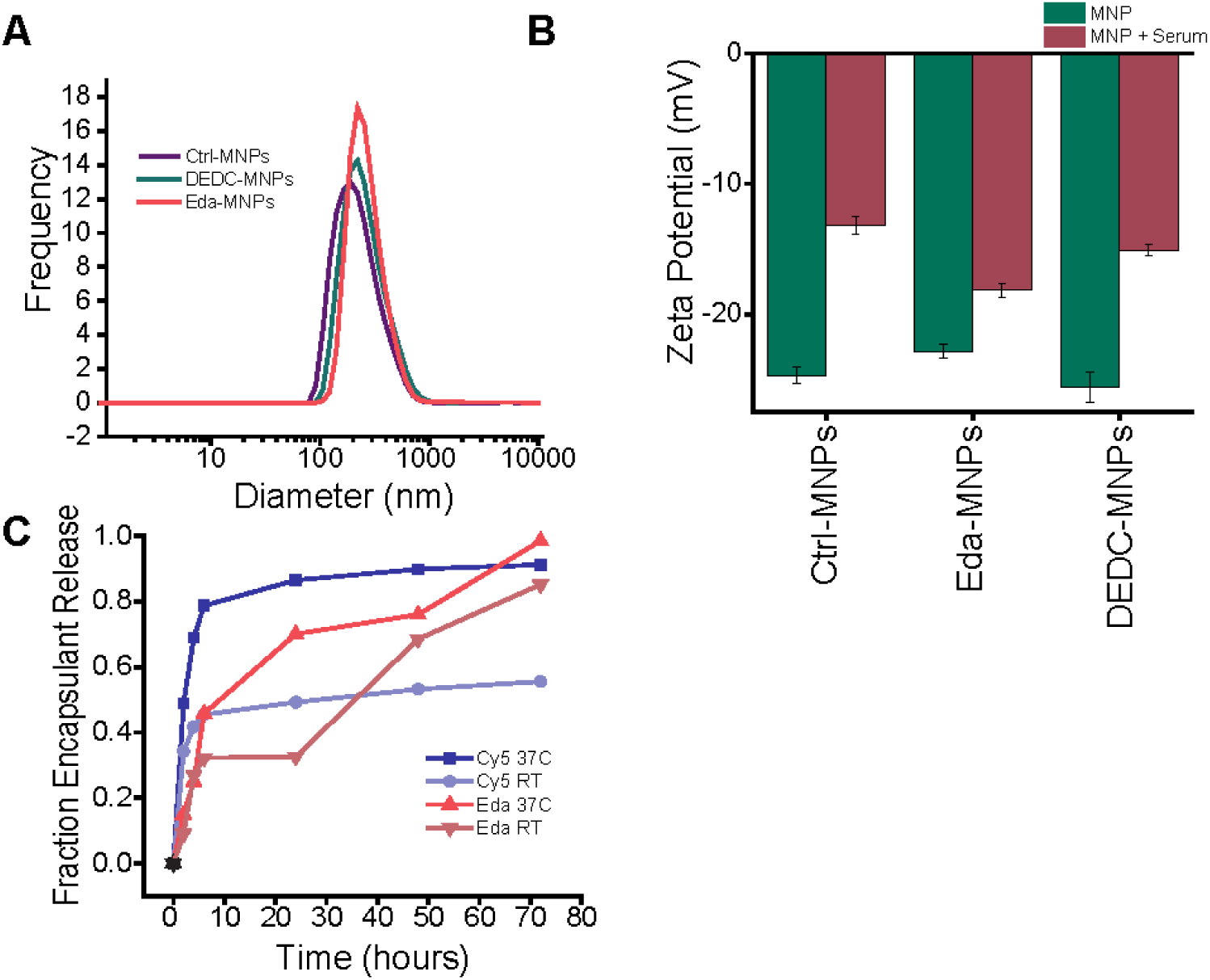
Mesoscale nanoparticle characterization. A) Hydrodynamic diameter distribution of Ctrl-MNPs, DEDC-MNPs, and Eda-MNPs used in all studies. B) ζ-potential of MNPs used in all studies in water versus after incubation with fetal bovine serum. C) Percent release of fluorescent dye (Cy5) or edaravone at room temperature (RT) or 37°C over time.

We performed in vitro cargo release assays to evaluate the period over which MNPs release the encapsulated dye or drug (Figure 1c). First, we found that Eda-MNPs contained 3.9 µg edaravone per 1 mg of particle while DEDC-MNPs contained 2.7 µg of dye per 1 mg of particle. Studies were performed in complete serum either at room temperature or 37°C, with measurements taken frequently up to 6 hours and daily to 72 hours. We found that DEDC-MNPs behaved similarly as before^17,18^, with approximately 40% of the cargo released in the first 6 hours before reaching a plateau at room temperature. At elevated temperature, the same pattern was followed, however about 80% of the cargo was release in the first 6 hours. Eda-MNPs exhibited similar release characteristics at both room temperature and 37°C (Figure 1c). However, only about 30% of edaravone was released at room temperature up to 6 hours and an additional 30% was released between 24 and 48 hours. Again, at elevated temperatures, the release rate was increased, though retaining the same pattern. It is interesting that while both cargoes exhibited a burst-release phase up to 6 hours, the remaining portion of bi-phasic release differed, with additional release for Eda-MNPs and only marginal release for DEDC-MNPs. We note that while release kinetics likely vary to due differences in edaravone and DEDC hydrophobicity and molecular weight^28^, these variations occur after 6 hours, a timepoint in which we would already expect particles to be localized to the kidneys^17,18^.

### MNPs Target the Kidneys in Tumor-Bearing Mice

We first aimed to investigate whether edaravone-loaded MNPs would continue to target the kidneys in tumor-bearing animals and avoid tumor localization that could result in the deactivation of cisplatin. The majority of prior work in nanoparticle-based drug delivery relied on the enhanced permeability and retention (EPR) effect to passively target tumors through leaky vasculature^29,30^. Were this to happen, reactive oxygen species scavengers would counteract the cytotoxic mechanism of cisplatin and mitigate cancer treatment^31^. This could potentially be a significant unintended effect of nanoparticle-based kidney targeting in patients or animals with advanced tumors.

To investigate whether MNP kidney localization indeed persists in the presence of tumors, we performed experiments in mice bearing human small cell lung cancer (SCLC) xenografts. We injected the luciferized NCI-H82 cell line^32^ subcutaneously into the flank of immunocompromised nude and NSG mice. We allowed these tumors to grow for approximately 1 month and injected 50 mg/kg DEDC-MNPs intravenously via the tail vein once tumors reached 500 – 750 mm^3^. Mice were sacrificed 24 hours later and fluorescence imaging of the tumor and organs was performed. We found that MNPs predominantly localized to the kidneys, with very little appearing in the tumors of either model (Figure 2a-b). In NSG mice, MNPs exhibited kidney-specific localization 6 – 18-fold greater than any other organ and 21-fold greater than the tumor (Figure 2a). In nude mice, we found this localization was 7 – 27-fold greater than any other organ measured and again 21-fold greater than the tumor (Figure 2b). This directly corresponds to prior work with a 50 mg/kg DEDC-MNP intravenous administration in healthy female hairless immunocompetent mice in which we observed 5-27 fold greater kidney targeting than other organs^17^. These data suggest that, even in the presence of quickly-growing xenograft tumors^25^, MNPs exhibit persistent renal localization which is not affected by the tumor or possible EPR effects.

**Figure 2.**
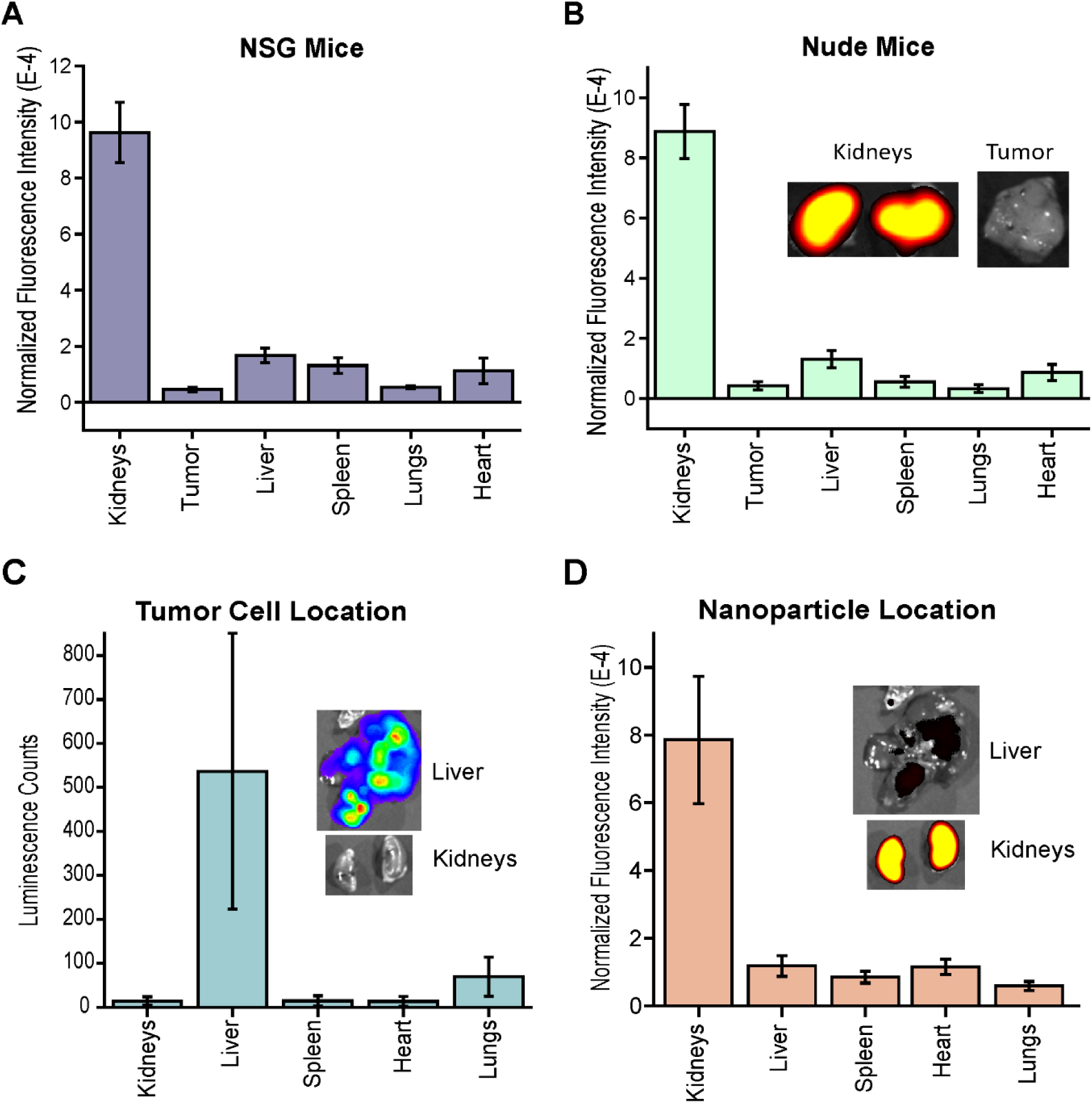
MNP biodistribution in tumor models. A) Normalized fluorescence of each organ to measure biodistribution in a NCI-H82 flank tumor xenograft model in NSG mice. B) Normalized fluorescence of each organ to measure biodistribution in a NCI-H82 flank tumor xenograft model in athymic nude mice. Inset: representative fluorescent image of kidneys and tumor with ex/em 650/680 nm. C) Bioluminescence imaging of each organ in an orthotopic metastatic tumor model in which mice were injected intravenously with NCI-H82 SCLC cells and allowed to grow for one month. Inset: representative bioluminescence image of liver and kidneys. D) Normalized fluorescence in each organ upon injection of DEDC-MNPs in mice from panel C. Inset: representative fluorescence image of liver and kidneys with ex/em 650/680 nm. Bars represent mean ± SD.

To further investigate MNP localization in the presence of tumors, we used an SCLC experimental liver metastasis model. NCI-H82 cells were injected via the tail vein into NSG mice and allowed to grow for approximately one month. DEDC-MNPs were injected intravenously via the tail vein and the animal was sacrificed 24 hours later after luciferin injection. Fluorescence and bioluminescence analysis of each organ was performed, confirming the presence of tumor primarily in the liver with modest growth in the lungs (Figure 2c). We found that DEDC-MNPS localize to the kidneys 7 – 13-fold greater than any other organ, including the liver (7-fold less accumulation than the kidneys) with substantial tumor signal and the lungs (13-fold less accumulation than the kidneys) with minor tumor signal (Figure 2d). Combined with prior data, this experiment suggests that MNP kidney-targeted biodistribution is not altered by the presence of tumors in the body.

### Eda-MNPs are Efficacious Against CI-AKI

To further evaluate the potential clinical application of MNPs, we next sought to recapitulate a mouse model of CI-AKI in non-tumor-bearing mice. To do so, first we induced CI-AKI in 8 – 10 week male C57BL/6 mice via intraperitoneal injection of 25 mg/kg cisplatin following 18 hours of fasting/water deprivation. We then sought to characterize whether MNPs localize to the kidneys in mice following cisplatin administration. At 24 hours following intraperitoneal cisplatin administration, we injected 50 mg/kg DEDC-MNPs via intravenous injection. Then, 48 hours after MNP administration, we sacrificed the mice and removed organs for fluorescence measurement. We again found that DEDC-MNPs primarily localized to the kidneys in mice following cisplatin injection (Fig. 3a). MNPs selectively localized to the kidneys 11 – 21-fold greater than any other organ measured, which favorably compares to the 5 – 27-fold greater kidney targeting with a 50 mg/kg intravenous MNP dose in healthy mice from prior work^18^. Following studies with Eda-MNPs, we also performed immunohistochemistry (IHC) with an anti-PEG antibody to stain for MNPs in the kidneys^18^ (Figure 3b). In agreement with previous studies, we found that Eda-MNPs selectively accumulate in primarily renal proximal tubular epithelial cells^17,18^. We then proceeded to therapeutic evaluation studies confident that MNPs exhibited similar biodistribution in the presence of disease initiation.

**Figure 3.**
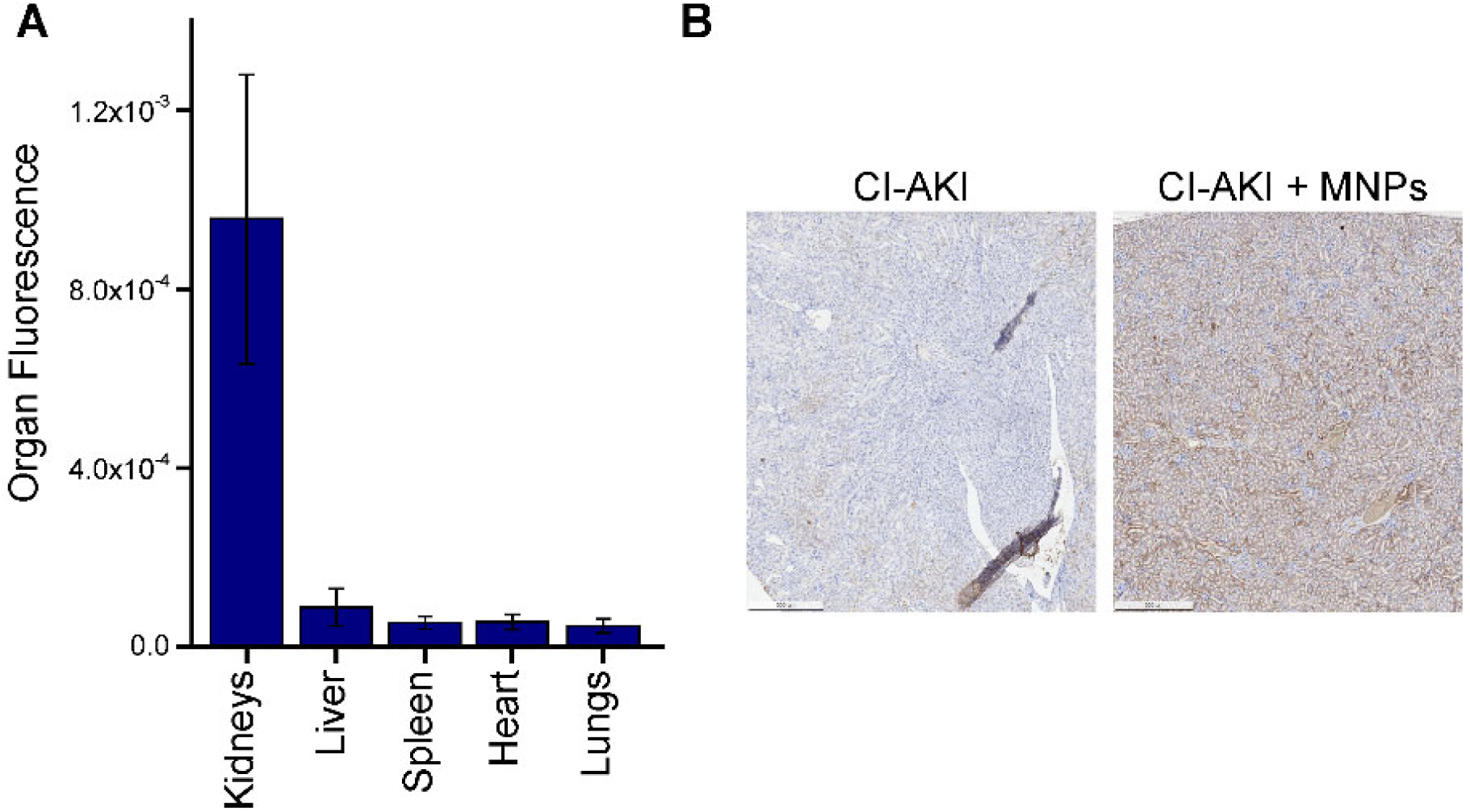
MNP biodistribution in mice administered cisplatin. A) Normalized organ fluorescence in each noted organ in mice injected with cisplatin and DEDC-MNPs 24 hours later, then sacrificed 48 hours after MNP injection. Bars represent mean ± SD. B) Immunohistochemistry with anti-PEG antibody staining for MNP accumulation in renal tissue.

To evaluate the therapeutic efficacy of Eda-MNPs against CI-AKI, we injected the particles 24 hours following cisplatin injection and injury initiation (Table 2). We performed additional control experiments in which we injected Ctrl-MNPs (empty) and two concentrations of free edaravone: 0.2 mg/kg to match that which was administered in MNPs and 30 mg/kg to match prior work with this drug in rat models of kidney injury^33^. These were compared against cisplatin-only and healthy control groups. We note that mice tolerated Eda-MNP, Ctrl-MNP, and the dose-matched edaravone IV injections well, however after a single high dose of free edaravone, mice exhibited acute belabored breathing and sluggishness from which they eventually recovered. At 48 hours after nanoparticle injection, and 72 hours after cisplatin administration, we sacrificed the animals and obtained serum and kidney tissue for analysis.

**Table 2.**
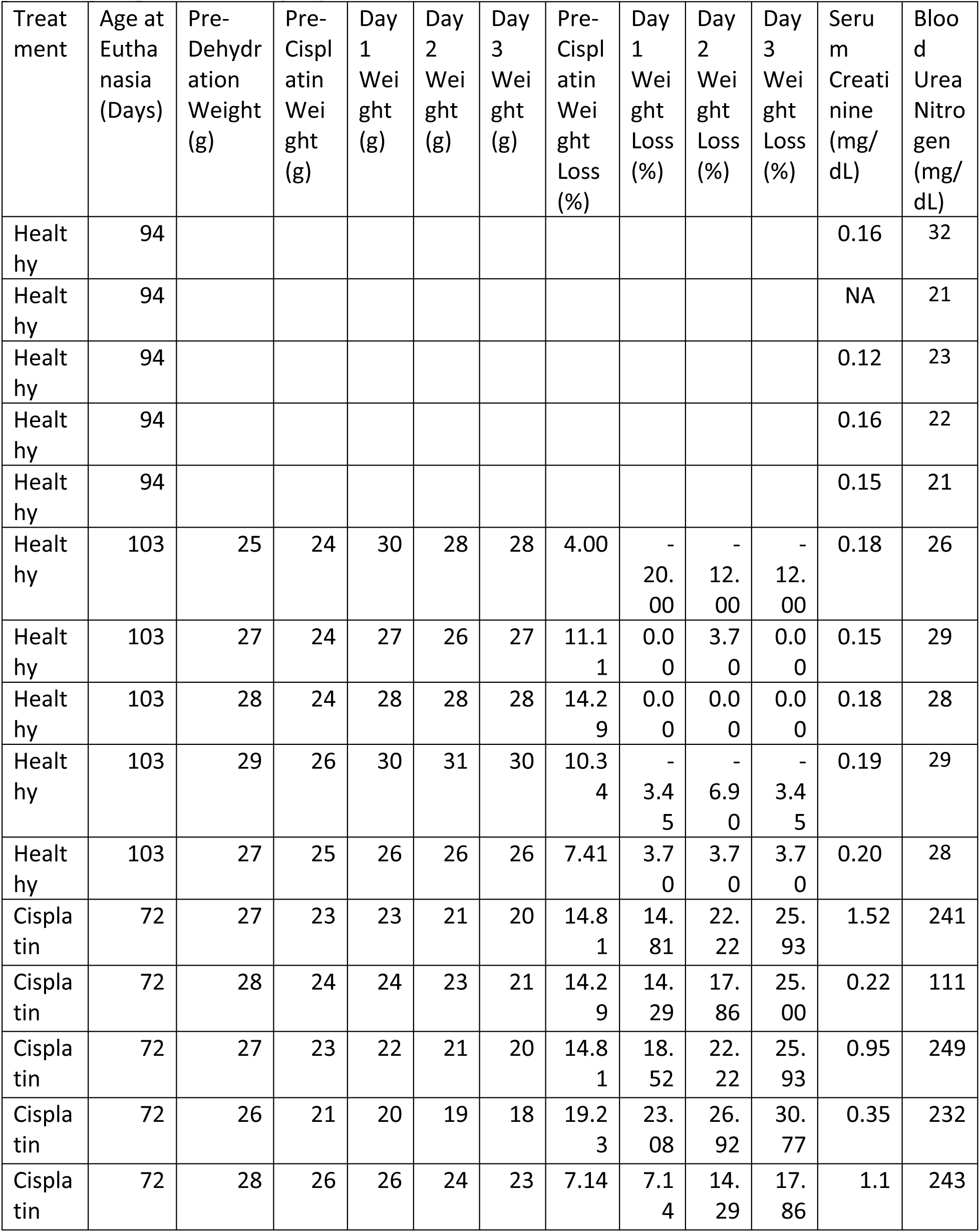
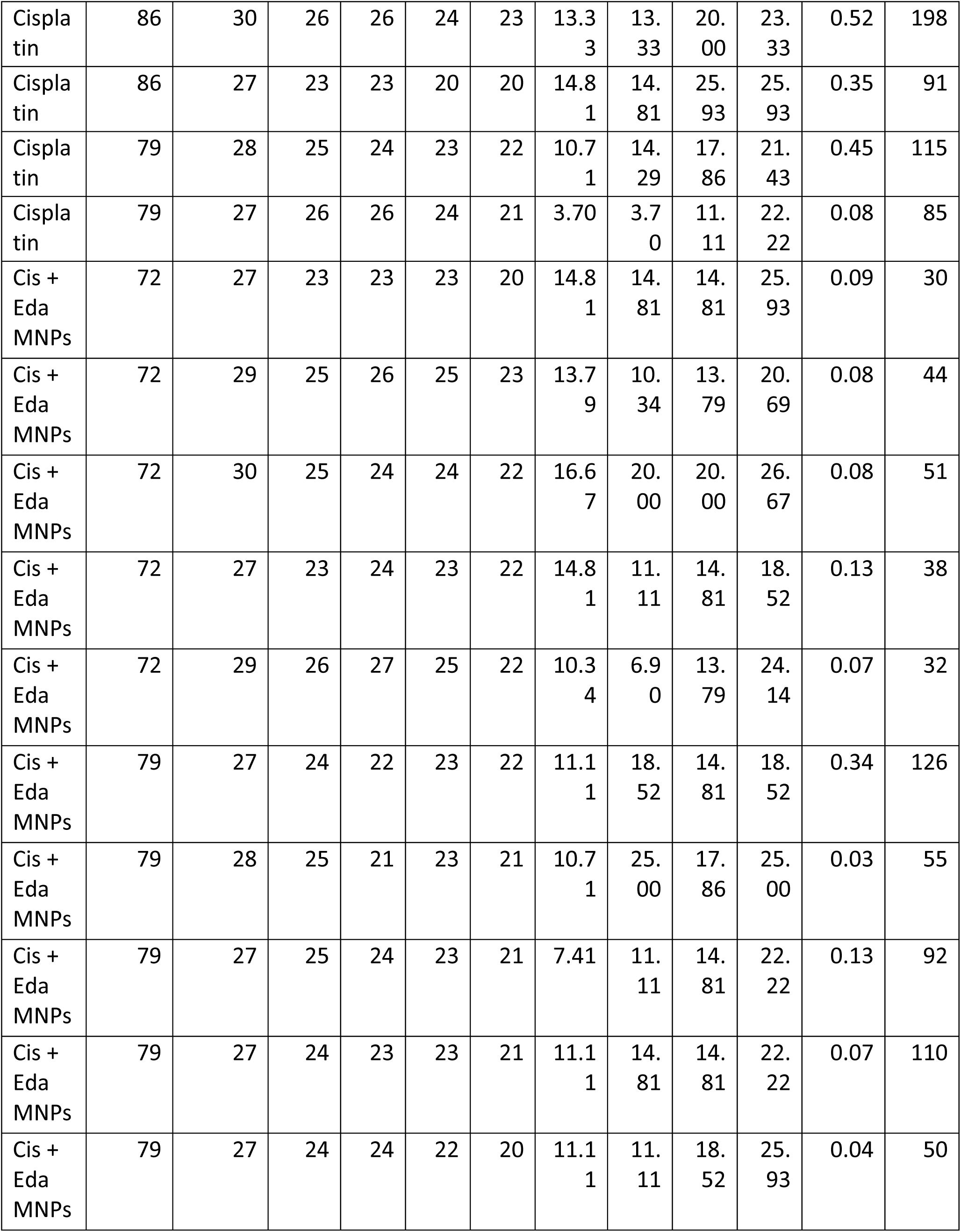
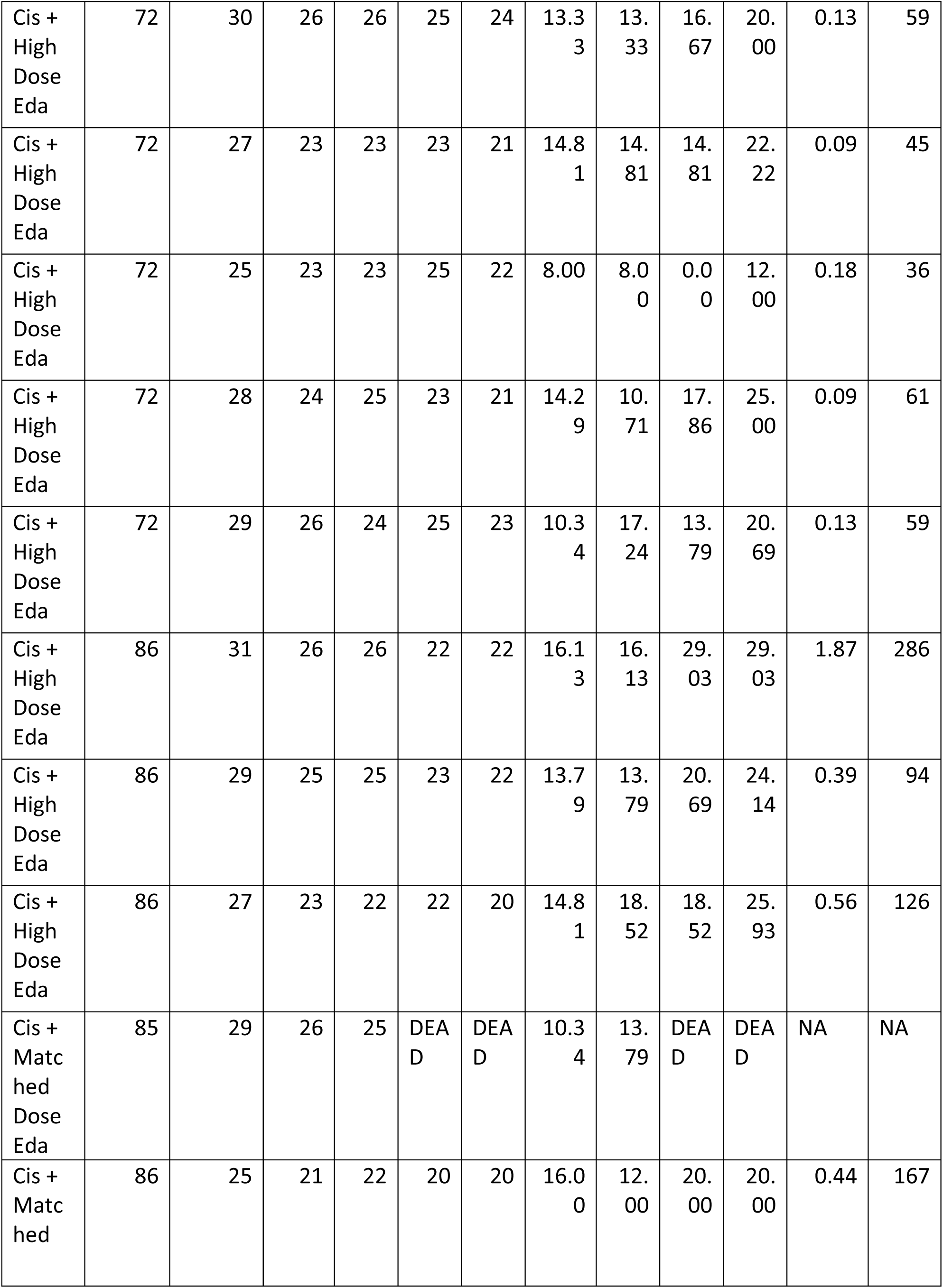
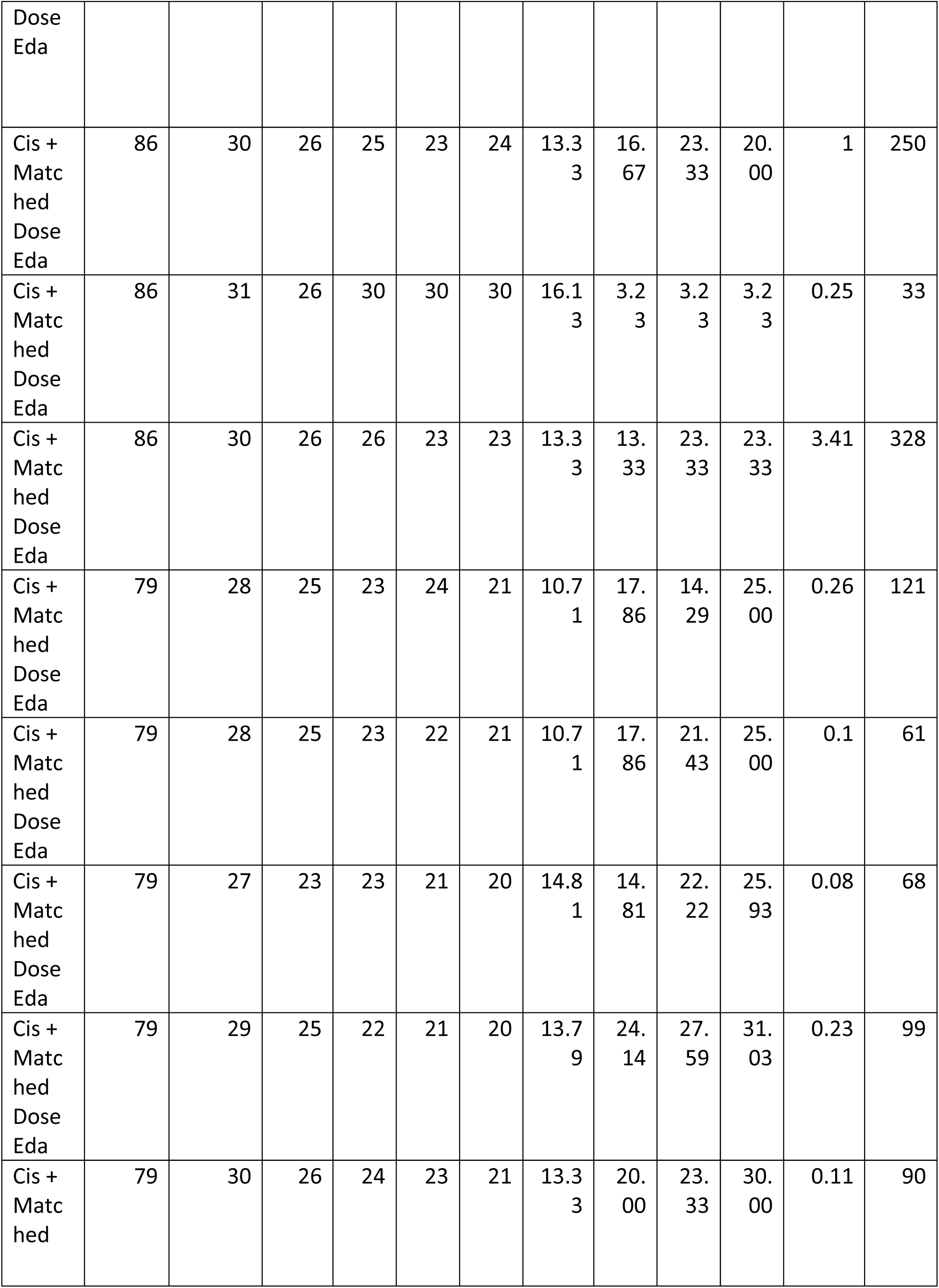
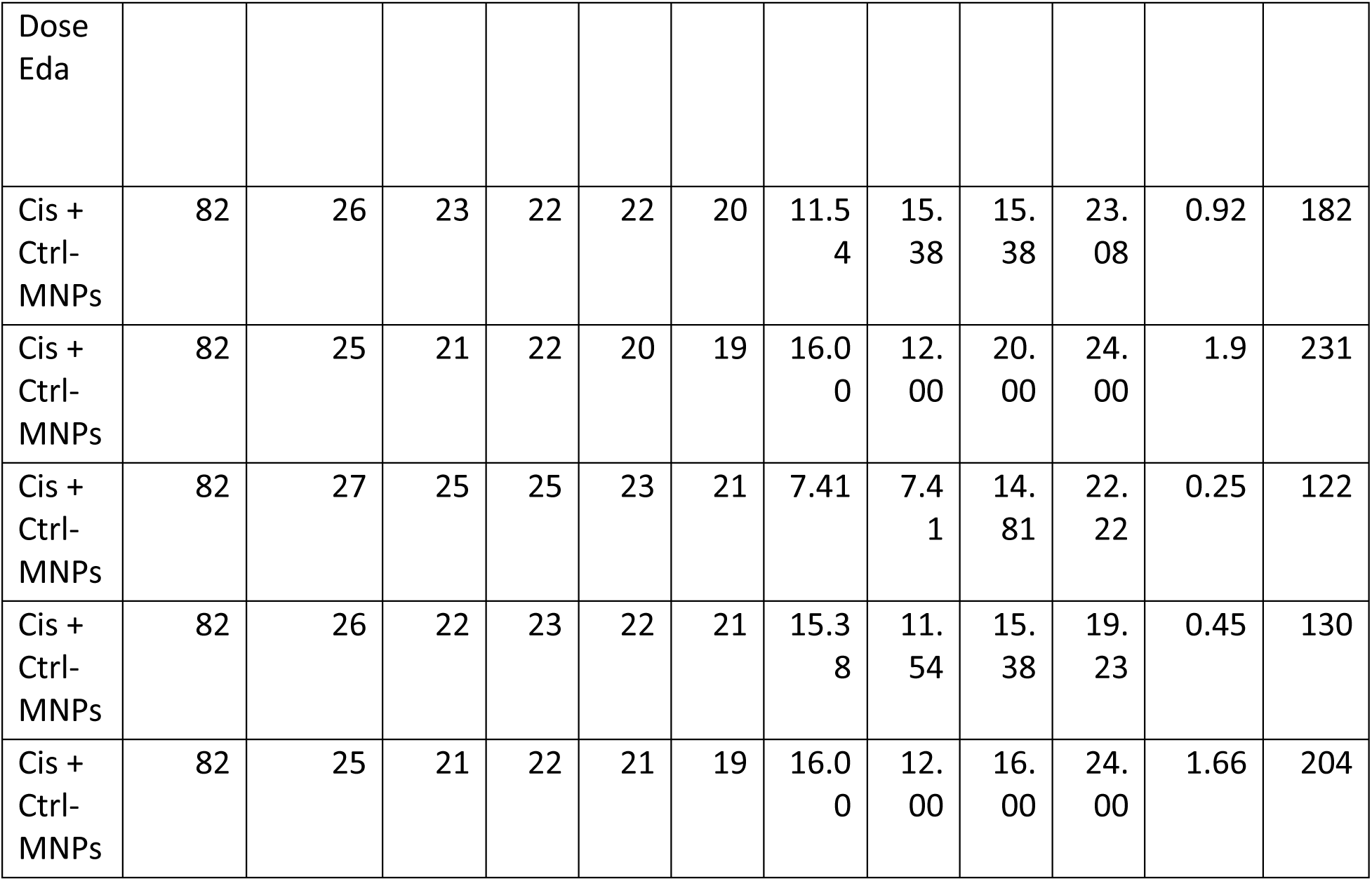
Individual mouse data from therapeutic efficacy studies. (note: a negative weight loss percentage represents a weight gain)

We found that mice with CI-AKI treated with Eda-MNPs exhibited kidney function and injury biomarkers similar to those in healthy control mice. We found that these mice had significantly lower serum creatinine (SCr) levels compared to untreated CI-AKI mice with levels at the baseline for healthy mice (Figure 4a; Table 2). We also found that the two free edaravone doses (dose-matched and high dose) did not have a significant effect on SCr levels, nor did the Ctrl-MNPs (empty). We also evaluated renal health through blood urea nitrogen (BUN) levels, finding the same evidence of therapeutic efficacy from Eda-MNPs (Figure 4b; Table 2), with only the high dose of free edaravone showing a moderate statistically-significant benefit. It is clear from these serum-based markers of renal function that Eda-MNPs exhibit substantial benefit to renal function following cisplatin insult.

**Figure 4.**
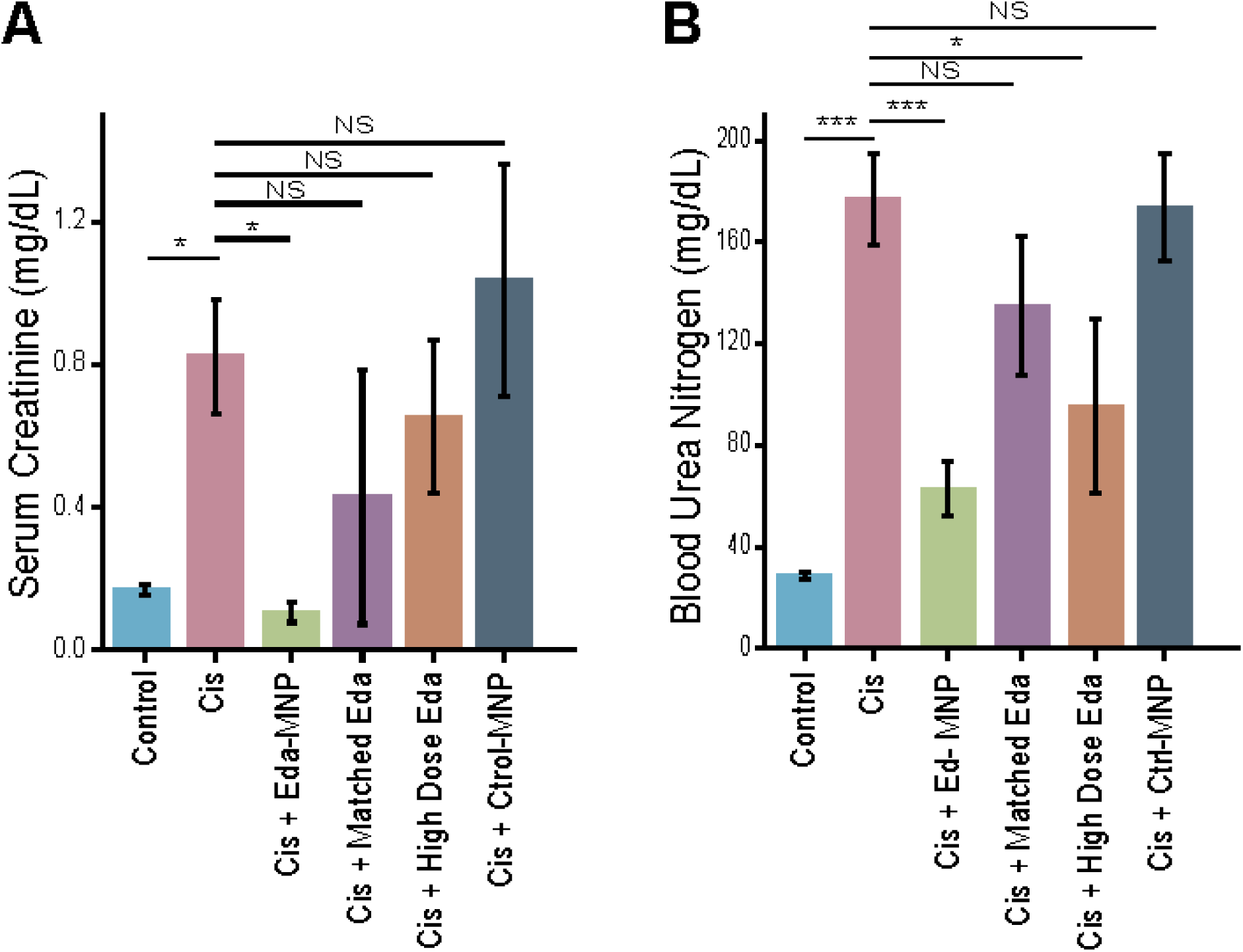
Therapeutic efficacy studies with Eda-MNPs and controls. A) Serum creatinine of mouse groups induced to have CI-AKI (Cis) and treated with Eda-MNPs or various controls. Bars represent the mean ± SEM. One-way ANOVA with Sidak’s posttest to compare to cisplatin control group. NS = p > 0.05; * = p < 0.05 B) Blood urea nitrogen (BUN) of the same mouse groups as in panel a. NS = p > 0.05; * = p < 0.05; * * * = p < 0.001.

To further investigate these findings, we performed histological analyses and IHC of renal tissue for each of the therapeutic and control groups. First, we evaluated hematoxylin and eosin (H&E) stains of each tissue, finding substantially reduced acute tubular necrosis, improved renal tubular architecture, and overall reduced evidence of AKI in mice treated with Eda-MNPs (Figure 5). Next, we performed TUNEL staining to identify DNA fragmentation and apoptosis, then IHC for p53 to identify loci of DNA damage and repair. Each of these clearly demonstrate a lack of apoptosis and DNA damage in mice treated with Eda-MNPs. We also performed IHC for the renal injury protein NGAL, finding substantially less injury in CI-AKI mice treated with Eda-MNPs. Finally, we performed IHC for nitrotyrosine to evaluate the pharmacodynamics of ROS scavenger therapy. Again, we found levels of oxidative stress that were comparable to healthy mice, and much less than CI-AKI mice or other controls. This data suggests that the ROS scavenger is having the intended effect at the site of injury, and in free edaravone controls little ROS scavenger is reaching the site of injury. Together, this data strongly suggests that Eda-MNPs have a significant therapeutic effect on renal health following CI-AKI.

**Figure 5.**
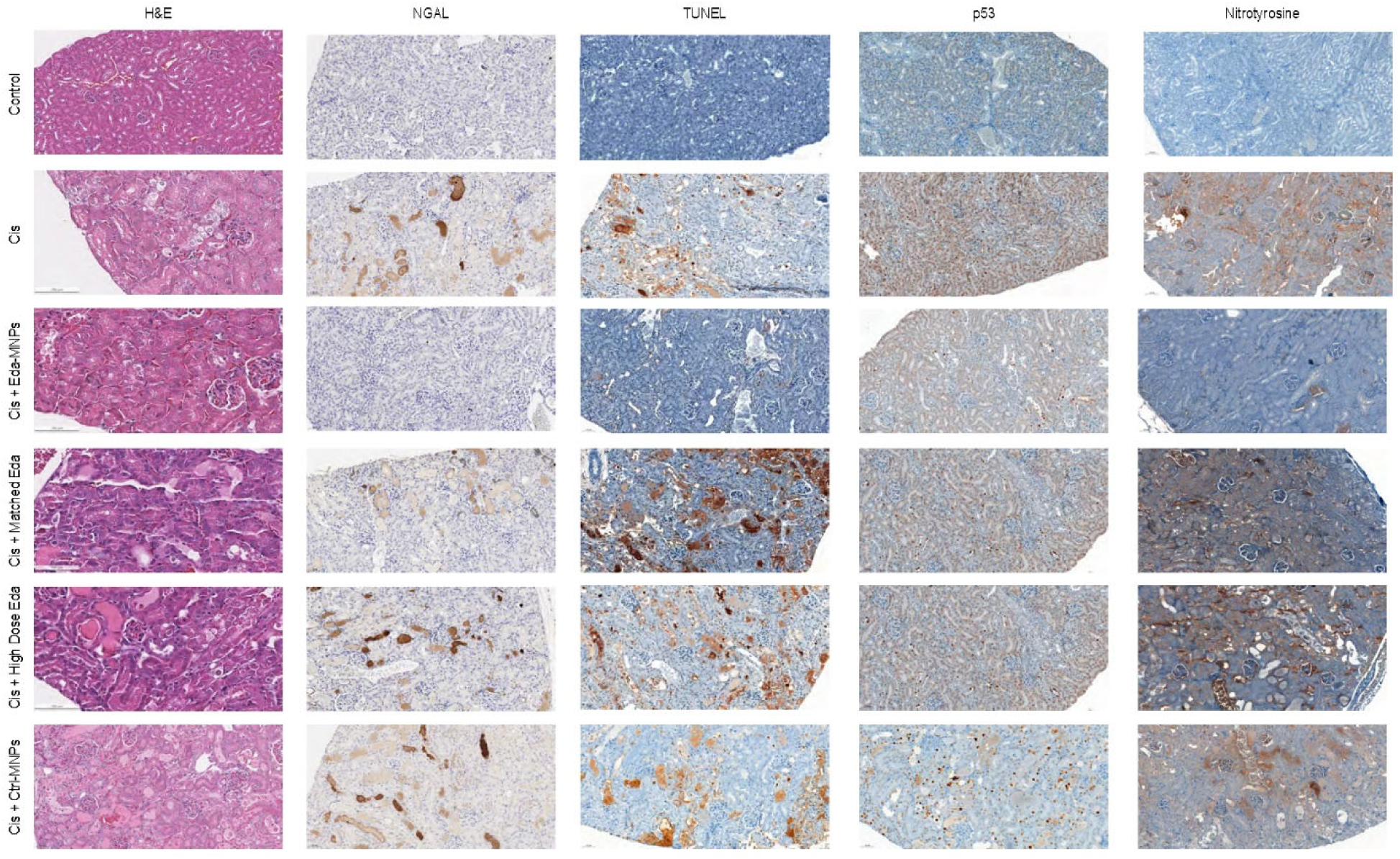
Representative histology of each mouse group in therapeutic efficacy studies. Tissues were stained with haemotoxylin and eosin (H&E) and IHC was performed for NGAL, TUNEL, p53, and nitrotyrosine.

## Conclusions

Tumor-specific targeting is a hallmark of passive nanoparticle localization and their clinical development^25^. However, it is essential that MNPs avoid this in order to maximize their clinical potential. We found that this indeed is the case, further suggesting that MNPs are suitable for clinical AKI therapy in patients with tumors. By administering edaravone via MNPs and not systemically, we would avoid interference with cancer treatment, as reactive oxygen species scavengers would do in cisplatin-treated patients^31^. By reducing renal complications, we also would allow the patient to stay on the front-line cisplatin therapy for longer periods of time at higher doses, extending the chemotherapy’s therapeutic window and effectiveness^34^. This advancement has clear utility in clinical cancer management and nephrology.

There have been few prior efforts to develop kidney-targeted drug carriers for renal disease, with even less focus on CI-AKI. Among these studies are DNA origami nanostructure approaches^35^, bacterial protein nanocage approaches^35^, and single-walled carbon nanotube^36^ approaches. These studies further validate the as-yet poorly-explored nanomedicine approach to the treatment of kidney disease^20^.

The MNP system here exhibits the most selective renal targeting compared to other organs as demonstrated in our prior work^17,18^ and in the studies reported here. As some of the other drug carrier systems are much smaller, there lies a possibility that they may accumulate in tumors via the EPR effect in animals or patients undergoing treatment for CI-AKI^25^. In this study, we found no safety issues with MNPs, which comports with our prior work in healthy mice^17^. However, the nanotube-based drug carrier also demonstrated safety in non-human primates^37,38^. We expect similar pre-clinical pharmacokinetic studies to further the possibility of MNP clinical translation.

In this work, we thoroughly investigated the renal targeting of mesoscale nanoparticles in mice bearing fast-growing flank xenografts or orthotopic metastatic tumor models, finding substantial kidney accumulation and virtually no tumor accumulation. We also found that drug-loaded MNPs exhibited substantial therapeutic potential in a murine model of CI-AKI. We found that MNPs target the kidneys in mice after cisplatin administration and successfully restored renal function in these animals with low doses of the free radical scavenger edaravone. These characteristics substantially increase the likelihood of translation to the clinic for the treatment or prevention of CI-AKI in patients with cancer. We expect this approach to ultimately improve the quality of those patients’ lives due to increased renal function after cisplatin therapy as well as a reduction in economic toll. We also expect this to allow those patients to stay on cisplatin as a front-line therapy, allowing increased survival from the underlying cancer.

## Materials and Methods

### MNP Formulation

Mesoscale nanoparticles were formulated from a di-block PLGA-PEG copolymer similarly to described in our prior work^17,18^. PLGA-PEG was synthesized via conjugation of carboxylic acid-terminated poly(lactic-*co*-glycolic acid) (PLGA) and heterobifunctional polyethylene glycol (NH2-PEG-COOH). PLGA (50:50; MW 38 – 54 kDa; 5 g; Aldrich, St. Louis, MO) was dissolved in 10 mL methylene chloride and mixed with N-hydroxysuccinimide (NHS; 135 mg; Aldrich) and 1-ethyl-3-(3-(dimethylamino)propyl)-carbodiimide (EDC; 230 mg; Aldrich) overnight. Activated PLGA-NHS was precipitated with cold ethyl ether and washed 3x with cold 50:50 ethyl ether:methanol before drying under vacuum. Dried PLGA-NHS (1g) was dissolved in chloroform with heterobifunctional PEG (250 mg; Nanocs, New York, NY) with 37 µL *N,N*-diisopropylethylamine and mixed overnight. PLGA-PEG was precipitated with cold methanol and washed 3x with the same before drying under vacuum. Conjugation was confirmed via ^1^H NMR as previously described^39^.

MNPs were formulated via nanoprecipitation. PLGA-PEG (100 mg) was dissolved in acetonitrile (2 mL) with either edaravone (100 mg; Santa Cruz Biotechnology, Dallas, TX) for Eda-MNPs, the Cy-5 mimic fluorescent dye 3,3’-diethylthiadicarbocyanine iodide (10 mg; Acros, Geel, Belgium) for DEDC-MNPs, or with no co-precipitate molecule for Ctrl-MNPs. This was added dropwise (100 µL / min) to purified water (4 mL) with Pluronic F-68 (100 µL; Life Technologies, Carlsbad, CA). Each batch of particles was stirred for 2 hours prior to centrifugation at 7356 RCF for 15 minutes prior to washing with purified water and centrifuging again. The recovered MNPs were suspended in a 2% sucrose solution and lyophilized until a fluffy powder was obtained.

### MNP Characterization

MNP batches were characterized to determine their size and charge. Size and polydispersity index (PDI) were measured via dynamic light scattering (DLS; Malvern, Worcestershire, UK) in 1x phosphate buffered saline (PBS) at 10 mg/mL. Surface charge was measured via electrophoretic light scattering (ELS; Malvern) in purified water at 1 mg/mL. The effect of serum on surface charge was determined by incubating each batch at 1 mg/mL in 100% fetal bovine serum (FBS) at room temperature for 30 minutes, centrifuging at 7356 RCF for 15 minutes, and resuspending the particle pellet in purified water for ELS. For each size and charge measurements, particle batches were measured three times and the mean ± standard deviation as reported.

Cargo loading and release were also characterized for Eda-MNPs and DEDC-MNPs. Cargo loading was determined by dissolving particles in acetonitrile, centrifuging at 31,000 RCF for 15 minutes to remove polymer, and performing UV-VIS spectrophotometry (Jasco, Easton, MD) on the supernatant. Cargo release was determined by suspending particles at 10 mg/mL PBS and incubating for 72 hours at 37°C or room temperature. At 2, 4, 6, 24, and 48 hours, as well as the final 72 hour timepoint, particles were centrifuged and the supernatant measured via UV-VIS spectrophotometry.

### MNP Targeting in Tumor Models

We first developed mouse tumor models to evaluate MNP biodistribution. To do so, we cultured the luciferized NCI-H82 small cell lung cancer (SCLC) cell line (a kind gift of S. Sharma, J. Lewis Lab, and E. Gardner, C. Rudin Lab, MSKCC)^32^. Cells were cultured in RPMI medium supplemented with 10% fetal bovine serum (FBS), 2 mM L-glutamine, 10 mM HEPES, 1 mM sodium pyruvate, 4.5 g/L glucose, 1.5 g/L sodium bicarbonate, and Primocin (MSKCC Media Preparation Core Facility).

We then implanted one million cells subcutaneously into the flank of four 8 – 10 week athymic nude male mice (Crl:NU(NCr)-*Foxn1*^*nu*^; Charles River, Wilmington, MA) after mixing with an equal volume of Matrigel matrix (Corning, Corning, NY) and allowed to grow 4-6 weeks. We also implanted the same number of cells subcutaneously into the flank of four male NSG mice (NOD-*scid* IL2Rgamma^null^; The Jackson Laboratory, Bar Harbor, ME) without Matrigel and followed the same procedures. DEDC-MNPs were injected intravenously at 50 mg/kg. Mice were sacrificed 24 hours later and we harvested the kidneys, liver, spleen, heart, and lungs as well as the tumor. Bioluminescence of all organs was imaged ex vivo after sacrifice via IVIS In Vivo Imaging System (Perking Elmer, Waltham, MA) to confirm tumor growth. Organs and the tumor were also subjected to IVIS fluorescence imaging with excitation/emission filters of 650/680 nm respectively. Each set of images was quantified via Living Imaging Software (Perkin Elmer) and normalized to a non-fluorescent control organ set.

We then injected intravenously via tail vein administration luciferized NCI-H82 cells into four 8-10 week NSG mice to establish orthotopic metastatic SCLC models. (Cells failed to grow following intravenous administration in athymic nude mice.) After 4-6 weeks and strong bioluminescence signal was obtained, we injected DEDC-MNPs intravenously at 50 mg/kg via the tail vein. Mice were sacrificed 24 hours later and organs were collected. Collected organs were imaged for bioluminescence and fluorescence as above. Each set of images was quantified via Living Imaging Software and normalized to a non-fluorescent control organ set.

### Cisplatin-Induced AKI Mouse Model

To next investigate the therapeutic efficacy of Eda-MNPs, we recapitulated a model of cisplatin-induced acute kidney injury as previously described^40,41^. Briefly, male 8 – 12-week C57BL/6 mice (Charles River, Troy, NY) were deprived of food and water for 18 hours prior to induction. We used male mice only as female mice are more resistant to renal injury^42-44^. Cisplatin (Sigma) was prepared for injection by dissolving at 1 mg/mL in sterile saline followed by a 30-minute incubation in a 37°C water bath to ensure dissolution while protecting from light. Mice were then injected intraperitoneally with 25 mg/kg, at which time food and water were returned. Mice were sacrificed 72 hours following cisplatin injection following a terminal retroorbital bleed. In addition to serum collection, kidneys were harvested and prepared for further study. Healthy control groups either underwent fasting and dehydration with no cisplatin injection or received food and water *ad libitum*.

### MNP Biodistribution in Cisplatin-Induced Mice

We first investigated MNP biodistribution in mice that were fasted, dehydrated, and injected with cisplatin as described above. In 3 mice, we injected DEDC-MNPs at 50 mg/kg via the tail vein 24 hours after cisplatin injection. 48 hours after MNP injection (and 72 hours after cisplatin injection), we harvested the kidneys, liver, spleen, heart, and lungs prior to IVIS imaging. Kidneys were formalin-fixed and prepared for anti-PEG immunohistochemistry (IHC) to confirm renal tubular distribution (HistoWiz Inc., Brooklyn, NY). Tissues were dehydrated and embedded in paraffin prior to obtaining 5 µm sections, placing on a glass slide, and deparaffinizing. Sections were heat-retrieved at pH 6 for 20 minutes and peroxide-blocked for 10 minutes. Slides were incubated with the anti-PEG antibody (1:400; ab94764; Abcam, Cambridge, UK) for 50 minutes and then incubated with a biotinylated rabbit anti-rat IgG for 8 minutes (AI4001; Vector Laboratories, Burlingame, CA). Slides were incubated with an anti-rabbit-conjugated horseradish peroxidase (HRP; Leica, Wetzlar, Germany) for 8 minutes and colorized with 3,3’-diaminobenzidine (DAB; Ventana Medical Systems, Tuscon, AZ). Hematoxylin (Ventana Medical Systems) was used to counterstain slides prior to Permount (Fisher Scientific, Waltham, MA) coverslipping. Digital slide images were obtained with a Leica AT2 slide scanner.

### Therapeutic Effect of Eda-MNPs in CI-AKI

We then sought to determine the therapeutic efficacy of Eda-MNPs in mice with CI-AKI. We investigated six total mouse groups (N = 5 – 10), including negative controls, positive controls, two unencapsulated drug groups, empty MNPs, and Eda-MNPs as the investigative group: 1) Healthy control mice; 2) CI-AKI; 3) CI-AKI + 50 mg/kg Eda-MNPs (intravenous in PBS); 4) CI-AKI + 0.2 mg/kg free edaravone (intravenous in PBS; matched dose); 5) CI-AKI + 30 mg/kg free edaravone (intravenous in PBS; high dose); and 6) CI-AKI + 50 mg/kg Ctrl-MNPs (intravenous in PBS). All treatments were performed 24 hours after cisplatin injection prior to sacrifice at 72 hours and organ formalin fixing and staining as above. Mice were weighed at the time of food and water removal, at cisplatin administration, and every 24 hours afterwards until sacrifice.

### Renal Function Analysis

At the time of animal sacrifice, blood was collected in serum preparation tubes (Becton Dickinson, Franklin Lake, NJ) and processed per manufacturer’s recommendation. Serum was analyzed for urea nitrogen and creatinine levels via colorimetric methods in the MSKCC Laboratory for Comparative Pathology. For CI-AKI therapeutic statistical analysis, a one-way ANOVA with Sidak’s posttest to compare to the cisplatin group was performed.

### Renal Histological Analysis

A portion of renal tissue from each group above was formalin-fixed and paraffin embedded as described above for histological analysis. Five parallel sections were obtained for each animal and subjected to the following treatments. 1) Hematoxylin and eosin (H&E) (MSKCC Molecular Cytology Core Facility or HistoWiz Inc.); 2) NGAL IHC (HistoWiz); 3) Nitrotyrosine IHC (MSKCC Core); 4) p53 IHC (MSKCC core); 5) TUNEL IHC (MSKCC Core).

## Acknowledgements

This work was supported in part by the NIH New Innovator Award (DP2-HD075698), the Cancer Center Support Grant (P30 CA008748), the National Science Foundation CAREER Award (1752506), the American Cancer Society Research Scholar Grant (GC230452), the Pershing Square Sohn Cancer Research Alliance, the Honorable Tina Brozman Foundation for Ovarian Cancer Research, the Expect Miracles Foundation - Financial Services Against Cancer, the Anna Fuller Fund, the Louis V. Gerstner Jr. Young Investigator’s Fund, the Frank A. Howard Scholars Program, Cycle for Survival, the Alan and Sandra Gerry Metastasis Research Initiative, Mr. William H. Goodwin and Mrs. Alice Goodwin and the Commonwealth Foundation for Cancer Research, the Experimental Therapeutics Center, the Imaging & Radiation Sciences Program, and the Center for Molecular Imaging and Nanotechnology of Memorial Sloan Kettering Cancer. R.M.W. was supported by the Ovarian Cancer Research Fund [Ann Schreiber Mentored Investigator Award 370463] and the American Heart Association Postdoctoral Fellowship (17POST33650043) and the City College of New York Grove School of Engineering. The authors would like to thank A. Klausner, J. Harvey, Y. Shamay, P. Jena, J. Kubala, R. Sridarhan, M. Manzari, and R. Skelton for helpful discussions related to this manuscript. Additional data related to this paper may be requested from the authors.

